# Immuno-Impaired Expression of Synaptophysin, GFAP and Nissl Substances in the Cerebral Cortex of Diabetic Wistar Rats; Evaluation of Andrographis Paniculata Effects

**DOI:** 10.1101/2025.11.22.689937

**Authors:** I.O. Onanuga, R.O. Folarin, E.T. Muniru, R.B. Ibrahim, M.I. Usman, A.I. Jegede, O.O. Azu, T.A. Obaya, M.O. Kehinde

## Abstract

**Background and aim:** Diabetic hyperglycemia is associated with severe complications, including neuropathy and cognitive impairment. This study examines the neuroprotective effects of *Andrographis paniculata* (*AP*) on the cerebrum cortex of alloxan-nicotinamide-induced diabetic male Wistar rats.

**Methods:** Thirty-five male Wistar rats were randomly divided into five groups (A–E), each with seven rats. Diabetic hyperglycemia was induced via a single intraperitoneal injection of nicotinamide (110 mg/kg) followed by alloxan (120 mg/kg). Treatment included *Andrographis paniculata*, metformin and a combination of both.

**Results:** Untreated diabetic hyperglycemia resulted in significant cerebral damage, indicated by weight loss, decreased brain weight, and neuronal degradation. *Andrographis paniculata* treatment provided partial neuroprotection. Metformin demonstrated significant neuroprotective effects by reducing hyperglycemia, preventing weight loss, and preserving neuronal structure. Combination therapy suggested potential synergistic effects, showing improvements in blood glucose, body weight, brain weight, cerebral morphology, and histochemistry. Increased synaptophysin expression and reduced astrocyte activation via GFAP expression were observed with combination therapy.

**Conclusion:** These findings support the therapeutic potential of *Andrographis paniculata*, alone or combined with metformin, in managing hyperglycemic neuropathies.

## Introduction

*Andrographis paniculata*, commonly known as the “king of bitters,” is a medicinal herb widely utilized in traditional medicine across Bangladesh, China, India, Indonesia, Malaysia, Pakistan, the Philippines, and Thailand (1). Indigenous to South Asian countries like India and Sri Lanka, it is now cultivated globally for its therapeutic properties (2). Numerous studies have highlighted the diverse pharmacological properties of *Andrographis paniculata*, which include antioxidant, antimicrobial, antidiabetic, anticancer, and antiviral effects (1–3).

Diabetes mellitus, defined by the World Health Organization (WHO) as a chronic metabolic disease, is characterized by elevated blood glucose levels that can lead to severe damage to vital organs, including the heart, blood vessels, kidneys, eyes, and nerves (4). This growing global health concern is accompanied by complications such as diabetic neuropathy, which is marked by nerve and brain damage, resulting in pain, sensory loss, and cognitive impairment (5–7). While existing treatments for diabetes primarily focus on controlling blood glucose levels, there is an increasing interest in therapies that also address the neurological complications associated with diabetes.

Diabetic hyperglycemia, a hallmark of diabetes mellitus, is characterized by persistently elevated blood glucose levels resulting from inadequate insulin production, insulin resistance, or a combination of both (8,9). This condition contributes to the pathophysiology of diabetes and is a significant risk factor for various complications, such as cardiovascular diseases, renal impairment, retinopathy, and neuropathy (5,6). Chronic hyperglycemia has a profound impact on the nervous system, leading to diabetic neuropathy and increased susceptibility to cerebrovascular damage, both of which are associated with cognitive decline and brain structural abnormalities (10,11).

In the brain, diabetic hyperglycemia induces oxidative stress, inflammation, and mitochondrial dysfunction, all of which contribute to neuronal apoptosis and impaired neuroplasticity (6,11). These mechanisms not only result in progressive cerebral atrophy but also exacerbate the risk of neurodegenerative conditions such as Alzheimer’s disease (12,13). The management of hyperglycemia is therefore crucial not only to control blood sugar levels but also to mitigate its neurotoxic effects, which are particularly detrimental in the cerebral cortex and other regions associated with cognitive function.

Standard antidiabetic treatments, such as metformin, primarily aim to lower blood glucose (1,2,9); however, they often fall short of addressing the neurological effects of diabetic hyperglycemia. This has led to a growing interest in adjunct therapies, like *Andrographis paniculata*, which may offer additional neuroprotective benefits. Given the combined need for blood glucose regulation and neuroprotection in diabetic patients, exploring treatments that simultaneously target hyperglycemia, and neurodegeneration is of increasing interest.

Both *Andrographis paniculata*, known for its anti-inflammatory and antioxidant properties, and metformin, a widely used antidiabetic drug, have demonstrated neuroprotective effects (14,15). However, the antidiabetic and combined effects of *A. paniculata* treatments on the histomorphology, histochemistry, and immunohistochemistry of the cerebrum in diabetic models remain unexplored. This study aims to evaluate the morphological and immunohistochemical impact of *Andrographis paniculata*, alone and in combination with metformin, on the cerebral cortex of alloxan-nicotinamide-induced diabetic male Wistar rats.

## Materials And Methods

### Drugs and Chemicals

Alloxan Monohydrate procured from (Sigma-Aldrich, St. Loius, MO, USA) which was of analytical grade was procured from UGOD lab supplies, Lagos, Nigeria. Nicotinamide and Metformin and Accu-Chek glucometer (Roche Diagnostics, Germany) were sourced from Yzee Pharmacy Ltd, Nigeria.

### Collection of Plant Material, Extraction and Screening

Fresh leaves of *Andrographis paniculata* were commercially purchased from a local herb store. Two kilograms of fresh *Andrographis paniculata* plant material was cleaned with tap water and air-dried in the dark at room temperature. The dried plant material was pulverized into a fine homogenous powder using an electrical blender (Binatone, United Kingdom). Two kilograms of the powdered plant was weighed and extracted using 70% ethanol through cold maceration. The solution was filtered through Whatman filter paper (Whatman International, United States), and the filtrate was concentrated to dryness using a rotary evaporator (Laborota 4000 - efficient, Germany) at 40 - 45°C. Phytochemical screening was conducted using standard precipitation and coloration tests. The identified major phytochemical constituents were tested as follows, Alkaloids, Flavonoids, Tannins, Saponins, Terpenoids, Phenols.

### Animal Care

Thirty-five (35) male Wistar rats, aged 6–8 weeks (150–210 g), were obtained from the Animal House, Babcock University, Ilishan-Remo, Ogun State, Nigeria. The animals were housed and managed at the Animal House of the Department of Anatomy, Olabisi Onabanjo University, Ago-Iwoye, Ogun State, Nigeria, under humane conditions according to the Principle of Laboratory Animal Care (Clark *et al*, 1997) and NIH Guide for the Care and Use of Laboratory Animals. The study protocol was approved by the Olabisi Onabanjo University Research Ethics Committee. The rats were housed in well-ventilated plastic cages having dimensions of 36 cm long x 24 cm wide x 15 cm high. They were maintained under standardized animal house conditions (temperature: 28–31°C; approximately 12hrs light/ 12hrs dark cycle per day) and were fed with standard rat chow (Premier Feed Mills, Nigeria) and given tap water *ad libitum* with saw dust used as beddings and other enrichment provided.

### Induction of Diabetes

Based on a prior pilot study, after eighteen (18) hours overnight fast, hyperglycemia was induced by single intraperitoneal dose of 110 mg/kg body weight of nicotinamide (Horbaach, USA) in physiological saline, and 15 minutes later, they were injected intraperitoneally with 120 mg/kg body weight of alloxan (Sigma-Aldrich, Chemical Company, Missouri, St. Louis, USA) dissolved in citrate buffer (pH 4.5) (16). The control groups received vehicle citrate buffer through the same route. Blood glucose levels were measured 72 hours post-induction using a glucometer (Accu-Chek, Roche Diagnostics, Germany). Rats with fasting blood glucose levels exceeding 100 mg/dL were considered hyperglycemic.

### Experimental Design

The animals were acclimatized for two weeks before the commencement of the experiment. The normoglycemic control (n = 7) comprises of seven animals while diabetic rats (n = 28) were divided into four treatment groups (B, C, D and E): Group A: received water as placebo (positive control); Group B: diabetic control (negative control); Group C: treated with 400 mg/kg of *AP* extract; Group D: treated with 1000 mg/kg of metformin; Group E: treated with 400 mg/kg of *Andrographis paniculata* extract and 1000 mg/kg of metformin. Administration was done daily via Oro-gastric gavage and the treatment lasted for four weeks.

### Blood Glucose, Food and Fluid Intake, and Weight Changes

Accu-Chek glucometer (Roche Diagnostics, Germany) was used to estimate the blood glucose of treated and control animals. Blood from the dorsal tail vein of the rats was used for measurement of capillary blood glucose concentrations on days 0, 7, 14, 21 and 28 (data not shown). Food and fluid intake was also monitored on a daily basis during the experimental duration (data not shown). Body weight (BW) changes of the animals were also recorded on the first day before the commencement of the treatment (the initial body weight), thereafter, weekly and on the last day shortly before animal sacrifice (final body weight). Organ weights were measured by an electronic balance (Mettle Toledo; Microsep (Pty) Ltd, Greifensee, Swizerland) post crainotomy. Values are expressed a gram for all weight measurements.

### Animal Sacrifice and Sample Collection

Four weeks after treatment, all animals were sacrificed by exposure to halothane for 3 min via a gas anesthetic chamber (100 mg/kg) at the end of the experimental period. Thereafter, blood samples were collected by cardiac puncture into pre-cooled heparinized tubes and serum bottles, placed on ice for 3 h and centrifuged in a desktop centrifuge model 90-1 (Jiangsu Zhangji Instruments Co., China) for 15 min at 3000 revolutions per minute. The serum was decanted into Eppendorf tubes and stored at –80°C for subsequent analysis. Brain tissues were excised, weighed and immediately fixed in 10% neutral buffered formalin. After proper fixation, tissues were dehydrated in a graded series of alcohol, cleared in xylene and embedded in paraffin wax using a cassette.

### Histopathological Examination of the Brain Tissues

For routine histological study, the paraffin-embedded brain tissues were sectioned at 5 μm thickness using an RWD S700A rotary microtome. The slides were deparaffinized in xylene and rehydrated in graded ethanol (100%, 90%, 80%, 70% and 50%) and rinsed in water. Slides were stained in hematoxylin for 5 min and rinsed with water, and counterstained in eosin for general assessment of the cerebral cortex morphology. The tissues were also stained with Cresyl Fast Violet (CFV) for visualization of Nissl bodies following Zhu et al. (2015)(17). The stained slides were then cover slipped using DPX mounting glue directly over the tissue sections. Thereafter, the slides were left overnight to dry for examination under a light microscope.

### Immunohistochemical Analysis

Separate slides were stained for each marker, Glial Fibrillary Acidic Protein (GFAP) and Synaptophysin, using the protocol of Shalaby et al. (2021)(11). The sections were blocked with goat serum to prevent non-specific binding. Primary antibodies used were mouse monoclonal GFAP (SC58766, Santa Cruz Biotechnology) and Synaptophysin (SC-17750, Santa Cruz Biotechnology), applied for 60 minutes each. After washing, the sections were incubated with a Biotinylated Secondary Antibody and developed using AEC as the chromogen, which stained positive cells brownish red.

The sections were examined using a binocular microscope, Leica ICC50 W, Germany (used to acquire the images). An independent histopathologist blinded to the treatment groups reported on the qualitative assessments of the slides. Image analysis for cell counts and staining intensity was performed using Image J software.

### Statistical Analysis

All data were expressed as mean + standard deviation (SD). Statistical comparison of the differences between the control and experimental groups was performed with GraphPad Prism software (version 8.00, GraphPad Software, San Diego, California, USA), using one-way analysis of variance (ANOVA), by the Turkey-Kramer multiple comparison test. A p-value of < 0.05 was considered significant.

## Results

### Average Weight Changes

As shown in TABLE 1, there was a highly significant difference in the initial and final body weights among the groups. When compared to the positive control group, rats treated with *Andrographis paniculata* and/or metformin displayed significant differences in their body weights, whereas no significant difference was observed in the body weights of the untreated diabetic group. Rats co-administered *Andrographis paniculata* and metformin exhibited a significant increase in final body weight relative to the positive control group. In contrast, untreated diabetic rats experienced a significant decrease in final body weight.

**TABLE 1.** STATISTICAL ANALYSIS OF BODY AND BRAIN WEIGHT.

| Groups | Initial BW (g) | Final BW (g) | BW diff (g) | % BW diff | Brain weight (g) | Relative brain weight |
| --- | --- | --- | --- | --- | --- | --- |
| Group A | 188.7±4.70 | 240.0±1.93 | 51.3 | 21.38 | 1.60± 0.04 | 0.007 |
| Group B | 204.7±3.47 | 142.4±5.35 * | -62.3 | -43.80 | 1.56± 0.01 | 0.011 |
| Group C | 221.3±2.23 * | 198.0±25.47 | -23.3 | -11.80 | 1.57± 0.10 | 0.008 |
| Group D | 247.3±1.65 * | 265.3±21.55 | 18 | 6.79 | 1.80± 0.03 | 0.007 |
| Group E | 275.0±6.97 * | 343.7±6.07 * | 68.7 | 19.99 | 1.91± 0.01 * | 0.006 |

Data is shown as mean ± SEM, $P \leq 0.05$ where $n$ is 7 for all groups. BW-Body Weight, DIFF- difference, %-percentage, \*= significant.

However, rats treated solely with *Andrographis paniculata* or metformin alone showed no significant differences in final body weight when compared to the positive control. Co-administration of *Andrographis paniculata* and metformin resulted in a significant weight gain, whereas the untreated diabetic rats showed a marked weight loss. Single treatments with either *Andrographis paniculata* or metformin alone did not significantly impact body weight.

Similarly, brain weight at sacrifice showed no significant differences among the groups relative to the positive control. A significant increase in brain weight was noted only in the group co-administered with *Andrographis paniculata* and metformin, as reported in TABLE 1.

### Serum Glucose Concentrations

TABLE 2 indicates significant differences in initial blood glucose levels (P= 0.0003). Compared to the positive control group, untreated diabetic rats and those treated solely with *Andrographis paniculata* showed significantly elevated initial blood glucose levels, while rats treated with metformin, either alone or in combination with *Andrographis paniculata*, displayed no significant differences in initial blood glucose.

**TABLE 2.**
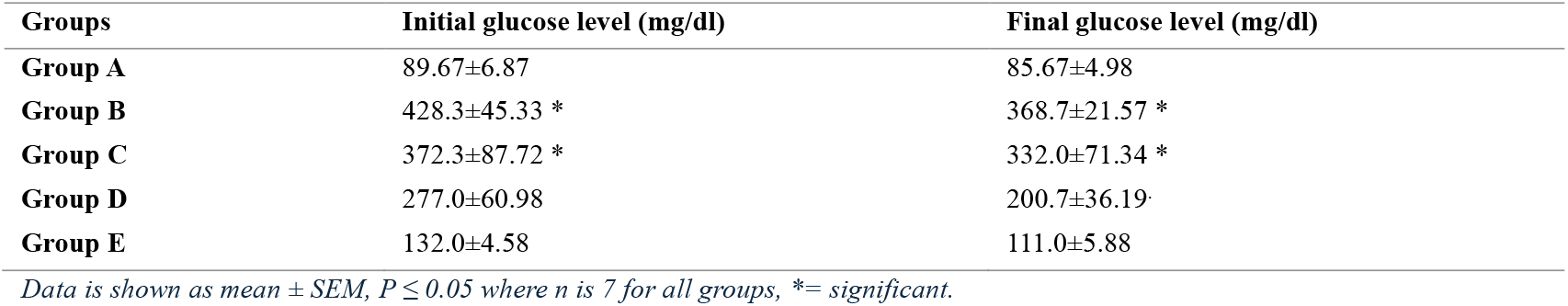
STATISTICAL ANALYSIS OF SERUM GLUCOSE BETWEEN DIFFERENT GROUPS.

Final blood glucose measurements revealed even more pronounced differences (P< 0.0001). Untreated diabetic rats and those treated with *Andrographis paniculata* alone displayed significantly elevated glucose levels compared to the positive control group. However, rats treated with metformin, both alone and in combination with *Andrographis paniculata*, showed no significant differences in final glucose levels when compared to the positive control. Furthermore, no significant differences were observed between initial and final blood glucose levels within any group (P=0.1655).

### Histological And Histochemical Evaluation of the Cerebral Cortex

Figure 1 (x100) and Figure 2 (x400) show the histoarchitecture of the cerebral cortex. The positive control, shown in Figures 1A and 2 A, exhibited normal cortical structure with regular cell distribution, neuronal morphology, and glial presence. The untreated diabetic group, shown in Figures 1B and 2B, displayed neuronal loss, cell swelling and extensive vacuolation.

**Figure 1.**
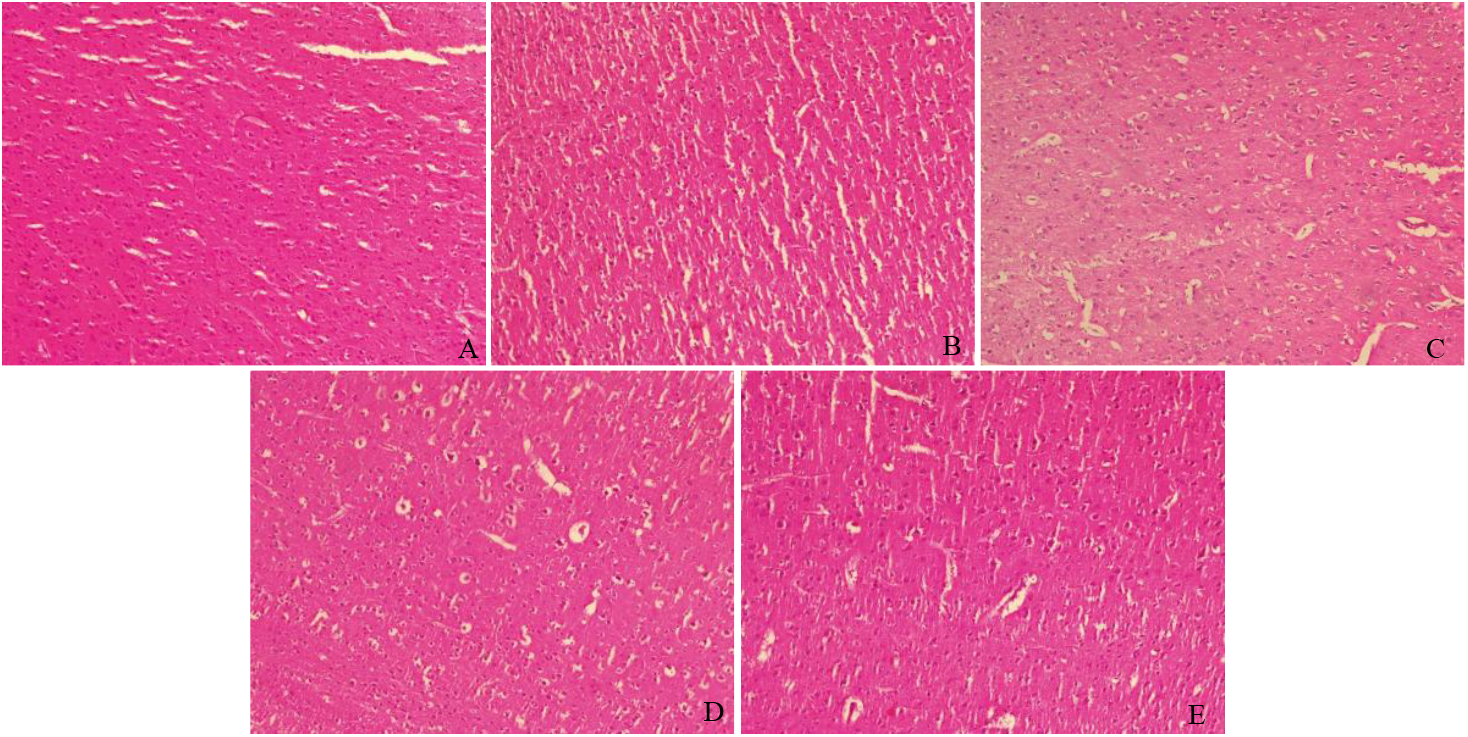
Photomicrographs of the cerebral cortex: [A] The positive control group, [B] The untreated diabetic group, [C] The Andrographis paniculata treated group, [D] The metformin treated group, [E] The metformin and Andrographis paniculata co-treated group. (H&E X100)

**Figure 2.**
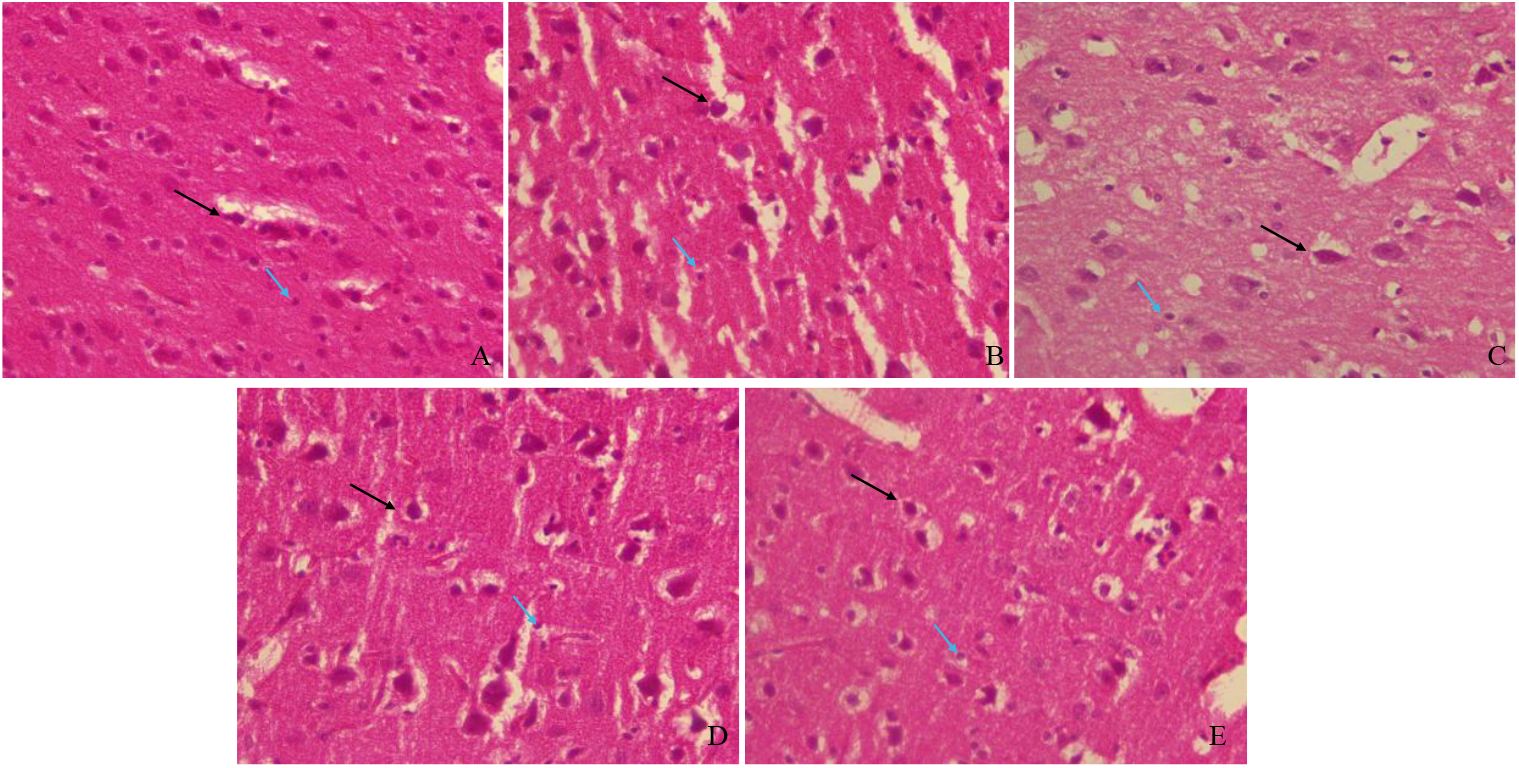
Photomicrographs of the cerebral cortex: [A] The positive control group, [B] The untreated diabetic group, [C] The Andrographis paniculata treated group, [D] The metformin treated group, [E] The metformin and Andrographis paniculata co-treated group. (H&E X400). Black arrows indicate pyramidal cells; blue arrows indicate neuroglial cells.

Rats treated with *Andrographis paniculata*, as depicted in Figures 1C and 2C, exhibited generally normal neuronal shapes, but with marked vacuolations and some signs of neuropathology. In the metformin-treated group (Figures 1D and 2D), neuronal shapes were largely intact with minimal vacuolation. The combination treatment group (Figures 1E and 2E) showed mostly intact neuronal morphology with minimal vacuolation and neuropathological indicators.

As seen in Figure 3, the positive control group (Figure 3A) showed well-stained pyramidal cells with regular vacuolation. In contrast, the untreated diabetic group (Figure 3B) displayed poor staining with substantial vacuolation, indicating reduced Nissl substance. The *Andrographis paniculata* treated group (Figure 3C) showed deeper staining of pyramidal cells compared to the untreated diabetic group, while the metformin-treated group (Figure 3D) displayed intense staining and reduced vacuolation. The combination treatment group (Figure 3E) exhibited both deep staining and regular vacuolation, indicative of improved cell integrity.

**Figure 3.**
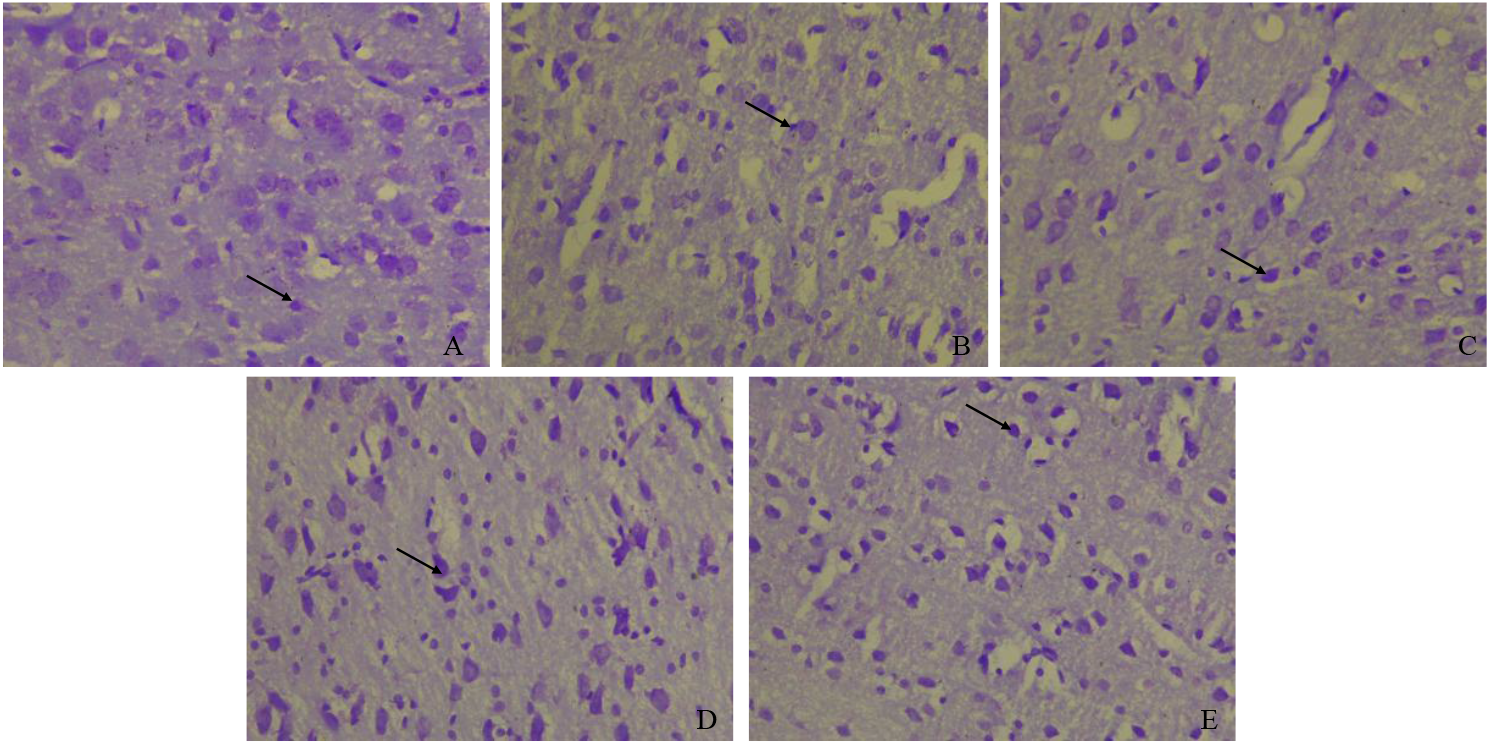
CFV staining of the Nissl bodies in the Purkinje layer of the cerebral cortex: [A] positive control group, [B] Untreated diabetic group, [C] Andrographis paniculata treated group, [D] Metformin treated group, [E] Metformin and Andrographis paniculata co-treated group. Black arrows indicate Nissl-stained cells.

### Immunohistochemical Evaluation of GFAP and Synaptophysin Expression

Figure 4 shows GFAP staining patterns. The positive control group demonstrated mild GFAP expression, indicative of minimal astrocyte activation. The untreated diabetic group, on the other hand, exhibited moderate GFAP expression with increased astrocyte activity. The group treated with *Andrographis paniculata* alone showed the highest GFAP expression, while the metformin-treated group exhibited the lowest GFAP staining. The combination treatment group showed mild GFAP expression, similar to that of the positive control, reflecting reduced astrocyte activation.

**Figure 4.**
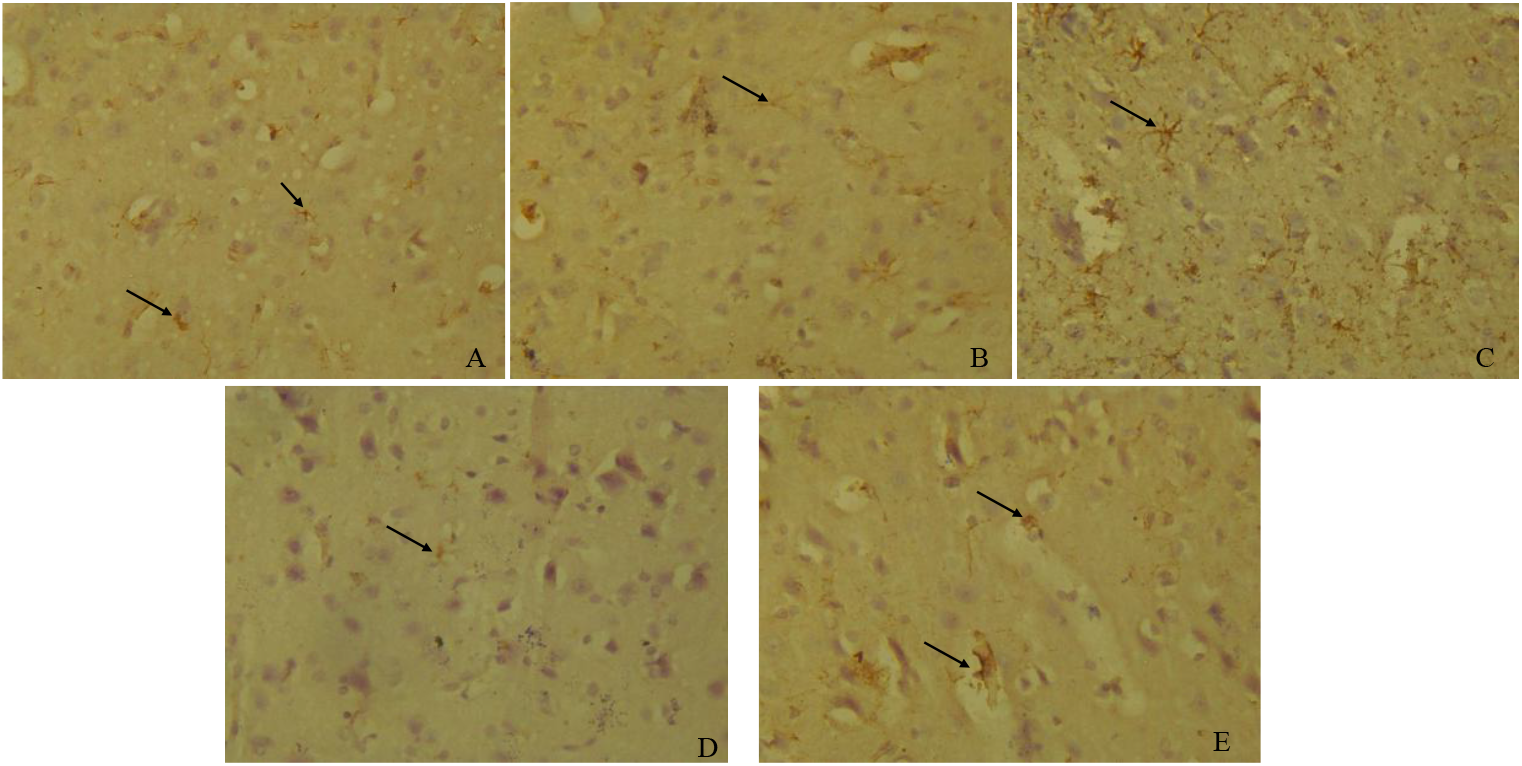
GFAP immunoassayed cerebral cortex section: [A] The positive control group showed mild GFAP expression in the astrocytes (arrows) of the Purkinje layer. [B] The diabetic untreated group displayed moderate GFAP expression in the astrocytes (arrows) of the Purkinje layer. [C] The group treated with Andrographis paniculata extracts alone displayed intense GFAP expression in the astrocytes (arrows) of Purkinje layer. [D] The group treated with metformin revealed weak GFAP expression in the astrocytes (arrows) of the Purkinje layer. [E] The group co-administrated with both Andrographis paniculata and metformin showed mild GFAP expression in the astrocytes (arrows) of Purkinje layer. (GFAP immunoassayed X400).

In Figure 5, synaptophysin expression across groups is shown. The positive control group displayed moderate synaptophysin expression, suggesting typical synaptic density. The untreated diabetic group, however, displayed weak synaptophysin staining, indicating reduced synapse density. The *Andrographis paniculata* treated group displayed moderate synaptophysin expression, comparable to the positive control. In the metformin-treated group, synaptophysin expression was mild, suggesting slight reductions in synaptic density. The combination treatment group exhibited strong synaptophysin expression, indicating a higher synapse density compared to other groups.

**Figure 5.**
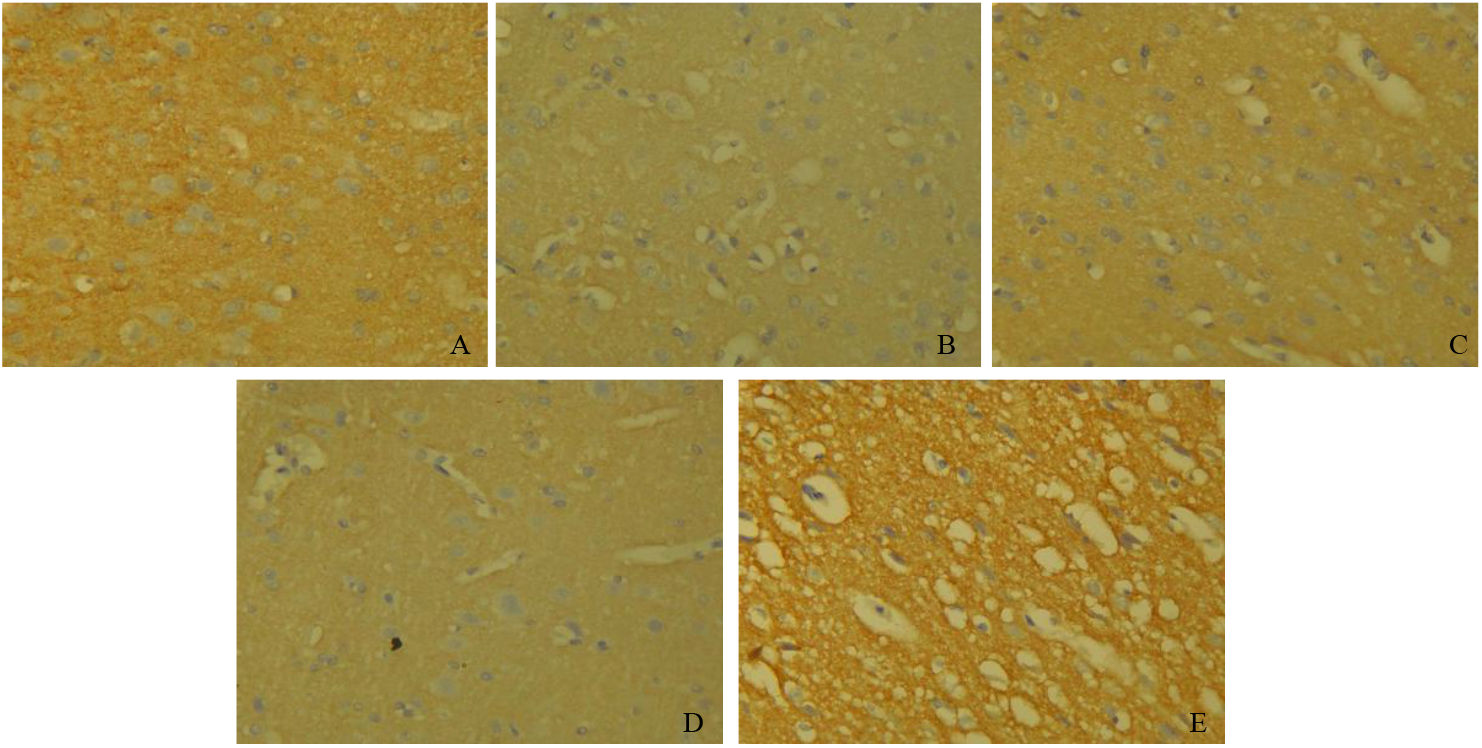
Synaptophysin immuno-stained cerebral cortex: [A] The positive control showed moderate synaptophysin expression. [B] The negative control showed weak synaptophysin expression. [C] The Andrographis paniculata treated group showed moderate synaptophysin expression. [D]The metformin treated group showed mild synaptophysin expression. [E] The Andrographis paniculata and metformin cotreated group showed strong synaptophysin expression.

### Histomorphometry Evaluation of Cerebral Cortex

The positive control group had the highest cell, and Nissl counts as reported in TABLE 3, the untreated diabetic group showed a significant reduction in both cell count and Nissl count. The Andrographis paniculata treated group had lower counts (cell and Nissl) but showed improvements over the untreated diabetic group. The metformin treated group showed substantial recovery in cell and Nissl counts. The Andrographis paniculata and metformin cotreated group showed lower cell count but higher Nissl count than the untreated diabetic group.

**TABLE 3.** STATISTICAL ANALYSIS OF THE HISTOMORPHOMETRY PARAMETERS.

| Parameters | Group A | Group B | Group C | Group D | Group E |
| --- | --- | --- | --- | --- | --- |
| Cell count | 183 | 136 | 95 | 129 | 117 |
| Nissl count | 163 | 57 | 67 | 110 | 85 |
| Cell/nissl ratio | 0.89 | 0.42 | 0.71 | 0.85 | 0.73 |
| Nuclear area | 1207±54.01 | 1362±79.87 | 1446±118.2 | 1476±115.7 | 1079±70.38 |
| Nuclear perimeter | 144.2±3.37 | 149.7±4.47 | 149±6.33 | 159.1±1.12 | 130.3±4 |

(Data is shown as mean ± SEM, where n is 7 for all groups).

The positive control had the highest cell/Nissl ratio while the untreated diabetic had the lowest cell/Nissl ratio. The Andrographis paniculata treated group and the metformin and Andrographis paniculata cotreated group showed comparable cell/Nissl ratios.

## Discussion

Diabetic hyperglycemia is known to contribute to long-term complications that affect multiple organs, including the eyes, kidneys, blood vessels, heart, and brain. These complications arise from a complex interplay of factors such as insulin resistance, oxidative stress, chronic inflammation, and metabolic disturbances. This study evaluated the effects of *Andrographis paniculata* on histomorphology and immunohistochemistry of the cerebral cortex in a diabetic hyperglycemic rat model.

The results showed that *Andrographis paniculata*, administered alone, led to a modest reduction in blood glucose levels compared to the untreated diabetic group, confirming prior findings on its antidiabetic properties (18). The observed stabilization in body weight and glucose levels in the treated groups underscores the therapeutic potential of *Andrographis paniculata* in diabetes. As expected, metformin, a well-established antidiabetic agent, significantly reduced blood glucose levels and helped maintain body weight compared to the positive control and untreated diabetic groups, which aligns with its use as a standard treatment for diabetes (19,20). Remarkably, the combination of *Andrographis paniculata* and metformin yielded the most significant reduction in blood glucose levels and an increase in body weight, suggesting a possible synergistic effect. This may be attributed to *Andrographis paniculata* anti-oxidative and anti-inflammatory actions, which could enhance metformin’s hypoglycemic effects, indicating a potential complementary mechanism of action.

The differences in sacrificial brain weight across groups suggest that both *Andrographis paniculata* and metformin, individually and in combination, may confer neuroprotection against diabetes-induced neurodegeneration. The significantly increased brain weight in the combination therapy group, relative to the untreated diabetic group, supports the notion that co-treatment may provide enhanced protection. This aligns with previous studies demonstrating that *Andrographis paniculata* and metformin individually possess neuroprotective properties, which are likely driven by antioxidative and anti-inflammatory mechanisms (14,21). Additionally, other studies have documented that both treatments promote cellular resilience and health in neural tissues (22,23).

The analysis of H&E and CFV staining revealed noticeable changes in neuronal morphology, vacuolation, and staining intensity across the treatment groups, reflecting varying levels of neurodegeneration and recovery. In the positive control, normal cortical histoarchitecture, cell distribution, and neuronal morphology were preserved. The untreated diabetic group exhibited severe neurodegenerative changes, such as swollen neurons, neurophagia, and vacuolations, consistent with studies that show hyperglycemia-induced neuronal damage through oxidative stress (8,24).

The *Andrographis paniculata*-treated group displayed some degree of neuroprotection, with improvement in neuronal shape and a reduction in neurodegenerative markers compared to the untreated diabetic group, though vacuolations were still present. This suggests partial neuroprotection, possibly due to the anti-inflammatory and antioxidant properties of *Andrographis paniculata* (23,25). The metformin-treated group exhibited a more pronounced restoration of neuronal morphology, with fewer vacuolations and degeneration markers, aligning with literature highlighting metformin’s neuroprotective potential (14). These effects may stem from metformin’s role in enhancing insulin sensitivity (7), reducing inflammation (26), and promoting neurogenesis (27).

The co-treated group showed the most significant improvements, with near-normal cell morphology and minimal signs of neurodegeneration, suggesting that *Andrographis paniculata* and metformin together provide a more comprehensive neuroprotective effect. The improved Nissl substance expression observed in CFV staining supports this conclusion, indicating restored neuronal function and communication.

Astrocyte activation, marked by increased GFAP expression, can signal neural stress or injury. In this study, moderate GFAP expressions were observed in the untreated diabetic group, consistent with findings on diabetes-induced astrocyte activation (6,11). *Andrographis paniculata* alone resulted in intense GFAP expression, which could be attributed to the study’s specific conditions or dosage, as this differs from some studies where *Andrographis paniculata* reduced astrocyte activation due to its anti-inflammatory properties (21,28). Conversely, the metformin-treated group showed weak GFAP expression, supporting its potential role in reducing glial activation and neuroinflammation (29). The combination group displayed mild GFAP expression, comparable to the positive control, suggesting that the combination may normalize astrocyte activation, which is beneficial in preventing neurodegeneration linked to excessive or insufficient astrocytic responses.

Synaptophysin serves as a marker for synaptic density, with reduced expression indicating synaptic loss commonly observed in diabetic neuropathy. In this study, the untreated diabetic group exhibited weak synaptophysin expression, indicating synaptic degradation, which aligns with studies that report synaptic vulnerability under hyperglycemic conditions (11,30). *Andrographis paniculata*-treated rats showed moderate synaptophysin expression, suggesting synaptic preservation, which aligns with its neuroprotective role. The metformin-treated group also showed mild synaptophysin expression, which may reflect metformin’s beneficial role in preserving synaptic integrity (27,29). Notably, the combination treatment led to the highest synaptophysin expression, suggesting a synergistic effect that may aid in synaptic preservation and recovery in diabetic neuropathy.

The differential outcomes observed between single and combination treatments of *Andrographis paniculata* and metformin emphasize the importance of exploring combination therapies for enhanced neuroprotection. The findings support the growing body of research indicating that combining natural compounds with pharmaceutical agents can mitigate diabetes-induced neurodegeneration and preserve neuronal health. This study highlights the therapeutic potential of *Andrographis paniculata* alongside standard antidiabetic drugs for combating neurodegenerative changes associated with diabetes.

## Conclusion

In conclusion, this study demonstrates that *Andrographis paniculata*, both individually and in combination with metformin, shows promise in ameliorating diabetic-induced neurodegeneration. The findings suggest that these treatments, particularly when combined, can mitigate diabetic neurodegeneration by preserving synaptic density, reducing astrocyte activation, and improving neuronal morphology. Further research is needed to elucidate the precise mechanisms underlying these effects and to determine the optimal dosages for maximizing neuroprotection in diabetic neuropathy.

## Ethical Approval

The study received ethical approval from the Olabisi Onabanjo University Research Ethics Committee.

